# Floating offshore wind farms cause Japanese horse mackerel to congregate

**DOI:** 10.1101/2025.03.17.643605

**Authors:** Shimpei Tsuchida, Riko Kato, Shoko Nishitsuji, Sayano Anzai, S. S. B. Azmi, Shogo Tanaka, Satoshi Masumi, Jun Uchida, Takashi Aoshima, Katsuya Hirasaka, Shingo Fujimoto, Kenichi Shimizu, Miyuki Hirose, Mitsuharu Yagi

## Abstract

Floating offshore wind farms (F-OWFs) are becoming key components of renewable energy production, yet their ecological impacts on marine ecosystems remain largely unexplored. Using environmental DNA (eDNA) analysis in the East China Sea, this study investigated the tendency for Japanese horse mackerel (*Trachurus japonicus*) to congregate beneath F-OWFs. Water samples were collected at stations near an F-OWF and control stations farther away at various depths and seasons. A total of 115 samples were analyzed, and eDNA of *T. japonicus* was detected in 83% of all samples. eDNA concentrations were significantly higher near an F-OWF (F-OWF stations) than at control stations. The highest recorded eDNA concentration reached 2,280 copies/L at an F-OWF station, whereas the maximum concentration at control stations was 783 copies/L. Seasonal variations were also observed, with lower concentrations in summer and higher concentrations from autumn to spring. Generalized linear model (GLM) analysis further revealed that wind turbines had a significant influence on eDNA concentration, whereas other environmental variables, such as water temperature and depth, were not significant. These findings suggest that F-OWFs may function as artificial reefs, providing habitat for commercially important fish and influencing fish distributions at both spatial and temporal scales. However, potential conflicts with fisheries due to spatial restrictions, displacement of fishery resources, and increased navigation costs necessitate further long-term ecological and socio-economic assessments. Integrating eDNA monitoring with traditional survey methods, such as acoustic surveys and ROV observations, is crucial for coexistence of adaptive offshore wind farm management and sustainable fisheries. Future research should also explore the long-term effects of F-OWFs on fish assemblages and biodiversity to support evidence-based decision-making for offshore energy development.

## INTRODUCTION

Construction of wind power facilities as hubs for clean energy generation has rapidly expanded worldwide. In recent years, many countries have begun exploiting renewable energy sources to reduce greenhouse gas emissions (Graabak et al., 2016), and commercial-scale wind power installations have increased significantly globally. According to the Global Wind Energy Council (GWEC), an additional 1,021 gigawatts (GW) of wind power capacity were installed in 2023, representing a 13% increase over the previous year (Global Wind Energy Council, 2024). There are two main types of wind power generation: offshore wind farms (OWFs) and land-based wind farms (LWFs). OWFs offer several advantages over LWFs. Specifically, wind speeds are generally higher and more consistent over the ocean compared to onshore locations, leading to greater potential power generation (Leung and Yang, 2012; Msigwa et al., 2022). Large-scale deployment of OWFs contributes to a stable power supply (Kazama, 2012). In 2023, OWFs experienced a global capacity increase of 10.9 GW, a 24% rise from the previous year (GWEC, 2024). This trend is expected to accelerate in coming years (Díaz and Guedes Soares, 2020).

Nonetheless, ecological risks associated with offshore wind farms (OWFs) have not been adequately assessed. It is well known that noise emissions and electromagnetic fields generated during transportation, construction, and operation of OWFs can impact marine environments (Kazama, 2012). Currently, bottom-fixed offshore wind farms (B-OWFs) are the predominant type, with monopile structures being the most commonly used. This design involves driving a large steel tube (pile) vertically into the seabed, upon which a wind turbine is mounted (Chen and Kim, 2022). Due to the piling process, monopile structures have significant environmental and biological impacts. Indeed, construction activities can induce behavioral changes in marine organisms (Thomsen et al., 2006), and extremely high noise emission levels have been reported during installation (Norro et al., 2013). However, some studies suggest that post-construction OWFs may have positive effects on marine life. Offshore structures attract fish, potentially functioning as artificial reefs (Amponsah et al., 2014; Wilson JC, 2007). Additionally, under the United Nations Convention on the Law of the Sea (UNCLOS), except for construction and maintenance purposes, vessels are prohibited from approaching within 500 m of OWFs (Bonsu et al., 2024; UNITED NATIONS, 1982; Reckhaus, 2022). As a result, waters adjacent to OWFs effectively function as no-fishing zones, offering protection for marine organisms (Hammar et al., 2016) and potentially benefiting commercial fish species (Bailey et al., 2014). Indeed, it has been reported that populations of the harbor porpoise (*Phocoena phocoena*), which declined during OWF construction, recovered to pre-construction levels after completion (Vallejo et al., 2017). Furthermore, post-construction surveys indicate a significant increase in fish abundance (Backer and Hostens, 2017; Lange et al., 2010). Despite these findings, most research to date has focused on B-OWFs, whereas ecological risk assessments of floating offshore wind farms (F-OWFs) remain scarce due to their relatively recent commercial deployment (Farr et al., 2021; Rezaei et al., 2023). Unlike monopile-based designs, F-OWFs employ hybrid spar-type structures, characterized by an elongated cylindrical floating foundation composed of a concrete lower section and a steel upper section, anchored to the seabed using three mooring chains (Sato and Matsunobu, 2021). This design allows for deployment in deeper waters, significantly expanding potential installation areas. Given the anticipated large-scale deployment of F-OWFs in offshore regions, understanding their impact on marine ecosystems is an urgent priority.

In this study, using environmental DNA (eDNA) technology, we investigated whether floating offshore wind farms (F-OWFs) affect distributions of Japanese horse mackerel (*Trachurus japonicus*). Our study site was located off the coast of the Goto Islands, Nagasaki, Japan, where Japan’s only F-OWF is located (Fig. 1). Due to the consistently strong winds and rough seas in this area, it is challenging to keep a research vessel stationed there for extended periods. Therefore, instead of conventional direct capture or visual surveys, we employed eDNA technology (Jerde et al., 2011), which provides a safer and more efficient method of data collection. Advances in quantification techniques and statistical modeling have enabled eDNA analysis to detect presence or absence of target species and to estimate their relative abundance (Fukaya et al., 2022; Hinz et al., 2022). The probability of species detection and eDNA concentration increases with higher species density (Hering et al., 2018; Yamaguchi, 2018). We assumed that if F-OWFs serve as fish aggregation sites, concentration of *T. japonicus* eDNA should be higher in areas near the F-OWF than in surrounding locations. *Trachurus japonicus* is a commercially valuable species (Igeta et al., 2023). Increasing numbers of F-OWF installations may reduce fishing grounds, potentially causing economic losses to regional fisheries (Methratta et al., 2020). Our findings contribute to understanding the impact of human activities on marine ecosystems and provide critical insights for balancing sustainable energy development with fisheries resource management.

**Fig. 1.**
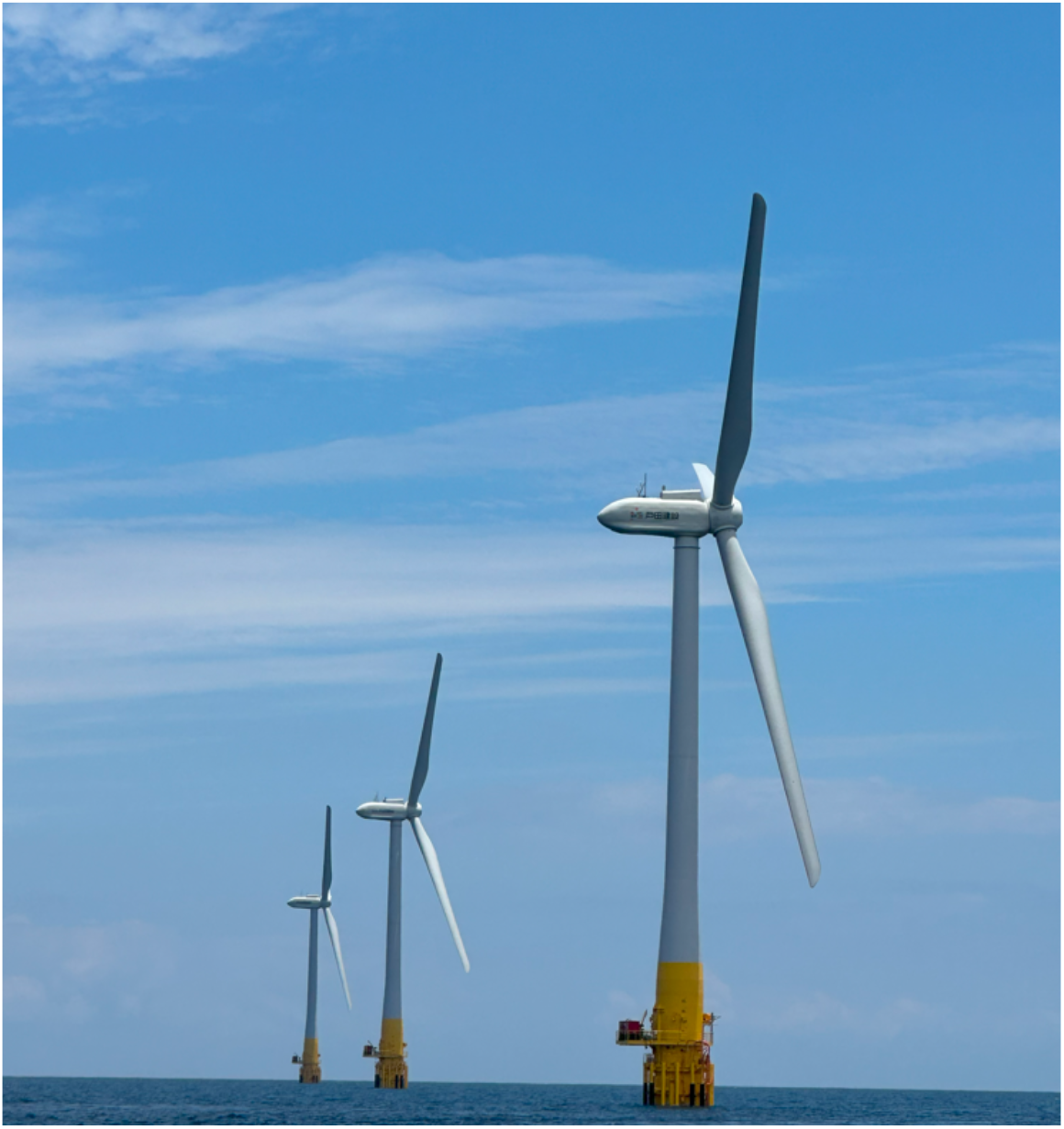
Floating offshore wind power facility (F-OWFs) taken in the experimental field of this study off the Goto Islands, Kyushu, Japan.

## MATERIALS AND METHODS

### Experimental design and water sampling

To evaluate the aggregation effect of F-OWFs on *T. japonicus* populations, we designed a field survey that incorporated both F-OWF and control sites. F-OWF sites included four stations (E1, E2, E3, and E4) near F-OWFs, approximately 5 km offshore from Fukue Island, Nagasaki, Japan. Control sites included four stations (E5, E6, E7, and E8), 4 nautical miles (7.4 km) south of the F-OWFs (Fig. 2). The F-OWFs in this region consist of one initial turbine, Haenkaze (FH), which was installed in 2015, followed by three additional turbines (F1, F2, and F3) in 2022 (Fig. 2). Each turbine has a rotor diameter of 80 m, a total height of 172 m (76 m below the surface and a hub height of 56 m), having a generation capacity of 2,100 kilowatts (kW). Latitude, longitude, and depth of each station and F-OWFs are provided in Supplementary Table 1.

**Fig. 2.**
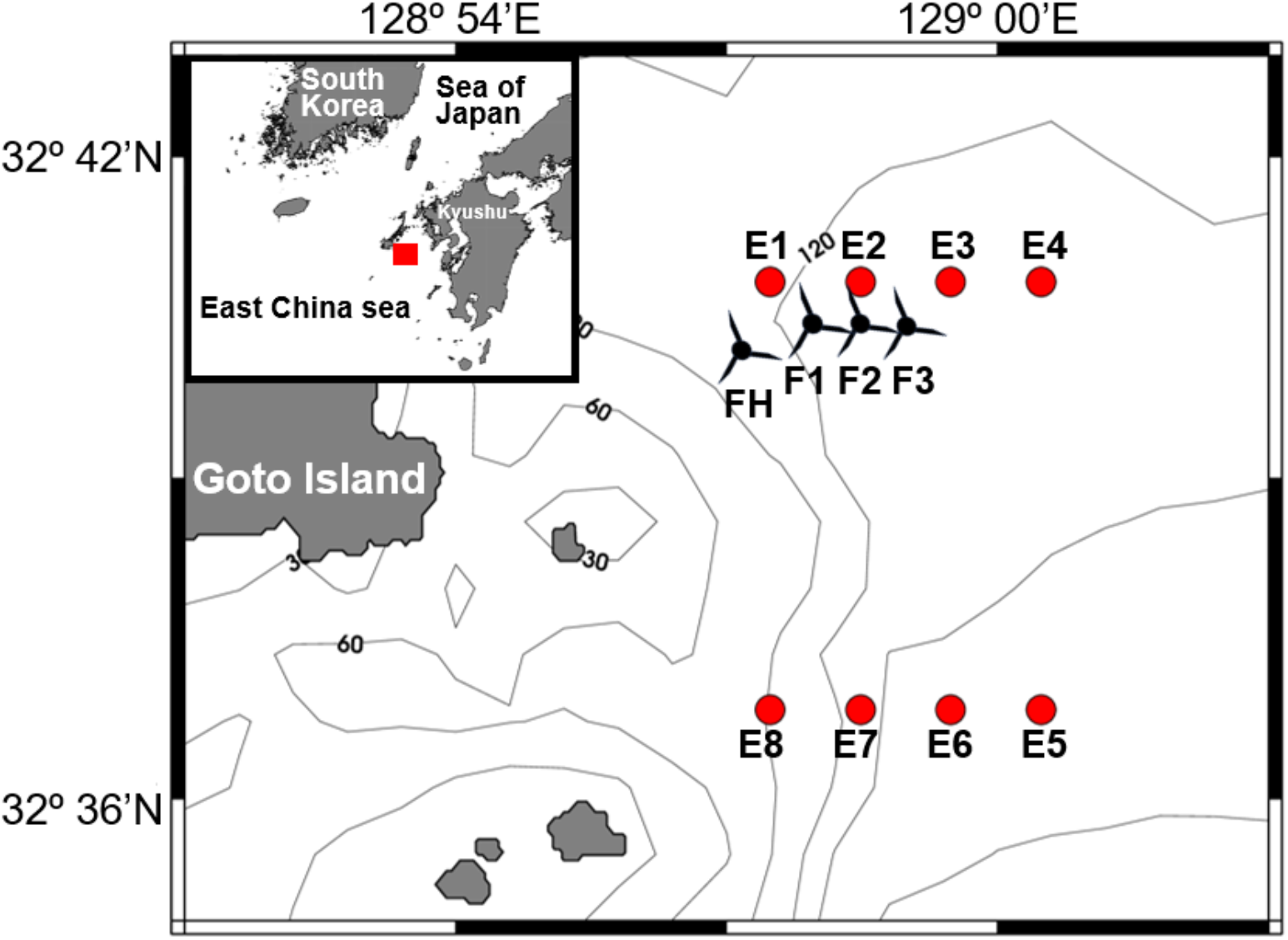
Bathymetric map showing the geographic locations of sampling stations (circles: E1~E8) in the East China Sea. E1~E4 is “F-OWF” and E5~E8 are “Control” stations. Floating offshore wind power facilities (F-OWFs: FH~F3) are also shown. The latitude, longitude, and depth of each station and F-OWFs are provided in Supplementary Table 1.

Water sampling was conducted during a research cruise of the *Kakuyo-maru* (155 tons), the training vessel of the Faculty of Fisheries, Nagasaki University, in April, June, August, October, and December 2023. At each station, water was collected from three depths: the surface layer (5 m), the middle layer (50 m), and the bottom layer (80–160 m). Sampling was performed using a CTD profiler (model number) installed on the training vessel, with a Niskin bottle to collect 3 L of sample water. The Niskin bottle and sampling hose were bleached with a chlorine-based detergent before sampling at each station to prevent contamination. Powder-free nitrile gloves were worn during handling. Using a CTD profiler for sampling significantly reduced the risk of contamination, which is a major challenge in eDNA analysis, while allowing precise water collection from designated depths. Seawater temperature was also measured at each sampling station as an oceanographic parameter. As a field blank, 3 L of artificial seawater were collected using the same method with a Niskin bottle. To inhibit DNA degradation, 3 mL of 10% benzalkonium chloride solution (final concentration: 0.01%) were immediately added to each sample after collection and samples were stored in a cool, dark place (Minamoto et al., 2021).

Within 48 h after collection, water samples were vacuum-filtered using an aspirator at the Fish and Ships Lab, Faculty of Fisheries, Nagasaki University. To prevent contamination during filtration, the filtration unit was covered with aluminum foil, and each device was decontaminated with 0.1% sodium hypochlorite after processing each sample. Filtration was performed using 47-mm glass fiber filters (Whatman GF/F No. 1825-047, pore size: 0.07 μm). Filters were wrapped in aluminum foil and stored at −20°C until eDNA extraction.

### DNA extraction and qPCR analysis

eDNA was extracted using a DNeasy Blood and Tissue Kit (Qiagen, Hilden, Germany) following a modified spin-column method based on the protocols of Ono et al. (2023, 2024) and Yamamoto et al. (2016). Blank samples were processed using the same extraction method. eDNA of *T. japonicus* was quantified using qPCR with the TaqMan probe method. Primers and probes followed Yamamoto et al. (2016) (forward primer: 5′-CAG ATA TCG CAA CCG CCT TT-3′, reverse primer: 5′-CCG ATG TGA AGG TAA ATG CAA A-3′, and probe: 5′-FAM-TAT GCA CGC CAA CGG CGC CT-TAMRA-3′). These primers specifically amplify a 127-bp fragment of the mitochondrial CytB gene of *T. japonicus*. The qPCR reaction mixture (total volume: 20 µL) consisted of 900 nM forward and reverse primers, 125 nM TaqMan probe, 2 × Environmental Master Mix (Thermo Fisher Scientific), AmpErase Uracil–Glycosylase (Thermo Fisher Scientific) and 2 µL of sample DNA. qPCR was performed using a QuantStudio 3 Real-Time PCR System (Thermo Fisher Scientific) under the following thermal cycling conditions: 50°C for 120 s, 95°C for 600 s, 55 cycles of 95°C for 15 s and 60°C for 60 s. To ensure quantitative accuracy, a standard curve was generated using a serial dilution of synthetic DNA fragments (1×10^1^, 1×10^2^, 1×10^3^, 1×10^5^, 1×10^7^, and 1×10^9^ copies). The sequence of the synthetic DNA fragment used was: 5′-CCT AGC TAT ACA CTA CAC CTC AGA TAT CGC AAC CGC CTT TAC ATC CGT AGC ACA CAT CTG CCG GGA CGT AAA CTA CGG CTG ACT TAT TCG CAA TAT GCA CGC CAA CGG CGC CTC CTT CTT TTT CAT TTG CAT TTA CCT TCA CAT CGG CCG AGG CCT TTA CTA CGG CT-3′. Each sample was analyzed in triplicate during each qPCR run. As a negative control, artificial seawater (2 µL) was analyzed via qPCR. Standard curves for all qPCR runs exhibited R^2^ values of 0.998–0.999, slopes of −3.67 to −3.56, and intercepts of 40.20–41.42. Based on these standard curves and Ct values of each sample, the mean copy number of the CytB gene fragment was calculated from three replicates per sample. No eDNA was detected in any of the negative controls from either field or laboratory experiments.

### Data analysis

Data normality and homogeneity of variance were assessed using the Shapiro-Wilk test and Bartlett’s tests, respectively. Given that data were not normally distributed, nonparametric tests were applied for statistical analysis. To compare mean eDNA concentrations between F-OWF and control stations, pooled across all sampling periods and layers, the Mann-Whitney U test was employed. Additionally, a two-way nonparametric ANOVA (Scheirer-Ray-Hare test) was conducted to examine effects of station (F-OWF vs. control), water layer (surface, middle, and bottom), and their interaction on eDNA concentrations. Furthermore, to analyze differences in monthly mean eDNA concentrations between F-OWF and control stations, a pairwise Wilcoxon rank-sum test with Bonferroni correction was performed following the Scheirer-Ray-Hare test to account for multiple comparisons.

To examine effects of environmental factors on eDNA concentration, a generalized linear model (GLM) analysis was employed. The response variable was the log-transformed eDNA concentration (log eDNA copies/L) (Sanchez et al., 2022; Maruyama et al., 2018). Explanatory variables included F-OWF presence or absence, water layer (m), and water temperature (°C) as environmental factors. F-OWF was coded as 1 and control as 0. The identity function was used as the link function, and a Gaussian distribution was applied. Variance inflation factor (VIF) values indicated no multicollinearity among the explanatory variables, as none exceeded 5 (F-OWF: 1.00, water layer: 1.76, water temperature: 1.75). The significance level was set at 0.05. All statistical analyses were conducted using R version 4.4.2.

## RESULTS

*Trachurus japonicus* eDNA was detected in 95 of 115 samples (83%). At F-OWF stations, 52 of 58 samples (90%) tested positive for eDNA, whereas at control stations, 43 of 57 samples (75%) tested positive. eDNA concentrations ranged from 8 to 2,280 copies/L at F-OWF stations and from 5 to 783 copies/L at control stations. When all monthly and water layer samples were pooled, the mean eDNA concentration was 322.6 copies/L ± 408.0 (standard deviation, S.D.) at F-OWF stations and 157.2 copies/L ± 174.6 (S.D.) at control stations, with a significantly higher concentration at F-OWF stations (*U* = 396, *n* = 115, *p* = 0.01), suggesting a potential attraction of F-OWFs for *T. japonicus* (Fig. 3).

**Fig. 3.**
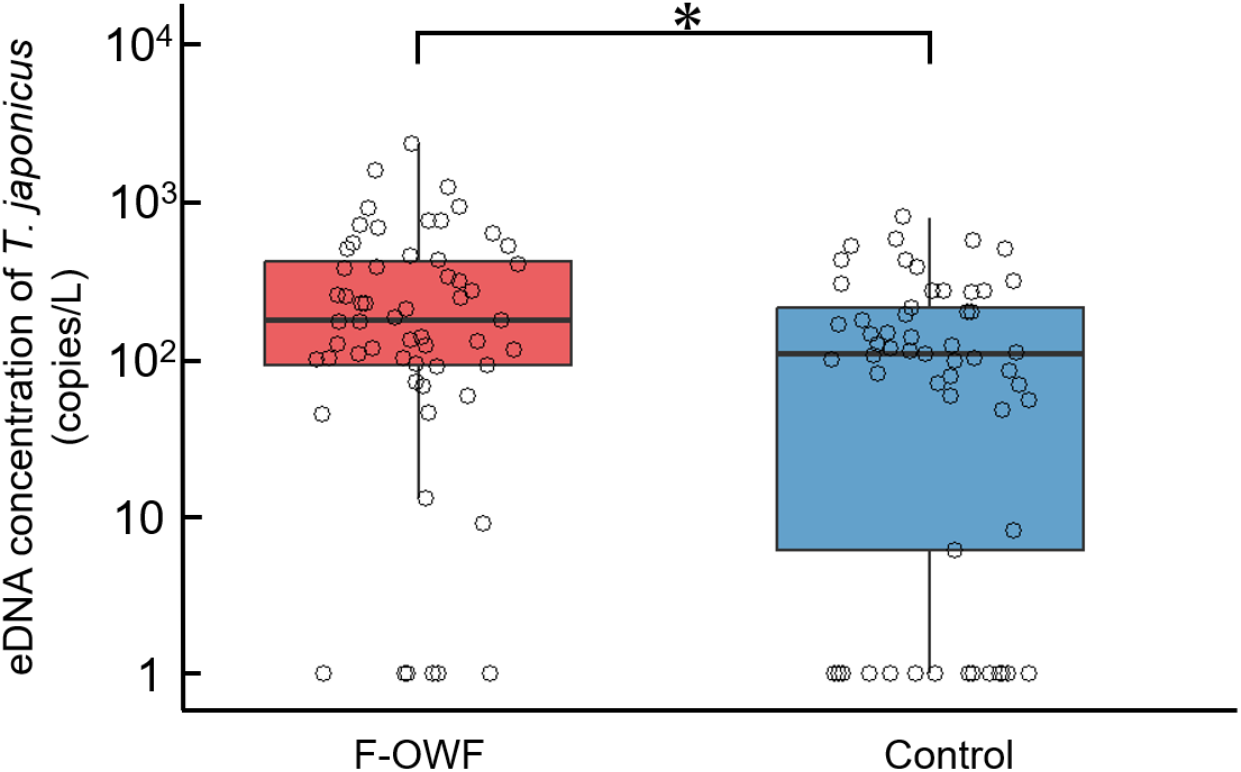
Comparison of *T. japonicus* eDNA concentrations between F-OWF stations and control stations, with data pooled across all sampling periods and water layers. The central line in the whisker plot represents the median, while the lower and upper edges of the box indicate the first (25%) and third quartiles (75%), respectively. Whiskers represent minimum and maximum values, and individual data points are displayed as a scatter plot. Statistical significance is indicated as * for *p* < 0.05.

Two-way nonparametric ANOVA was conducted to examine effects of sampling station (F-OWF or control) and water layers (surface, middle, bottom) on eDNA concentration. This analysis revealed that sampling stations had a significant positive effect on eDNA concentration (*p* = 0.01) (Table 1 and Fig. 4). In contrast, water layers and their interaction with sampling stations were not significant (*p* = 0.38 and *p* = 0.56, respectively) (Table 1 and Fig. 4), indicating that water layers do not influence eDNA concentration, regardless of the sampling station.

**Table 1.** Summary of two-way nonparametric ANOVA assessing effects of sampling station (F-OWF or control) and water layers (surface, middle, bottom) on eDNA concentration. Sampling stations had a significant effect on eDNA concentration. In contrast, water layers (surface, middle, bottom) and their interaction with station location did not significantly affect eDNA concentration. Statistical significance is indicated as “*” for *p* < 0.05.

| Factor | d.f | H statistic | p-value |
| --- | --- | --- | --- |
| Station<br>(Treatment/Control) | 1 | 1.96 | 0.01* |
| Layer<br>(Surface/Middle/Bottom) | 2 | 6.31 | 0.38 |
| Interaction<br>(Station × Layer) | 2 | 1.17 | 0.56 |

**Fig. 4.**
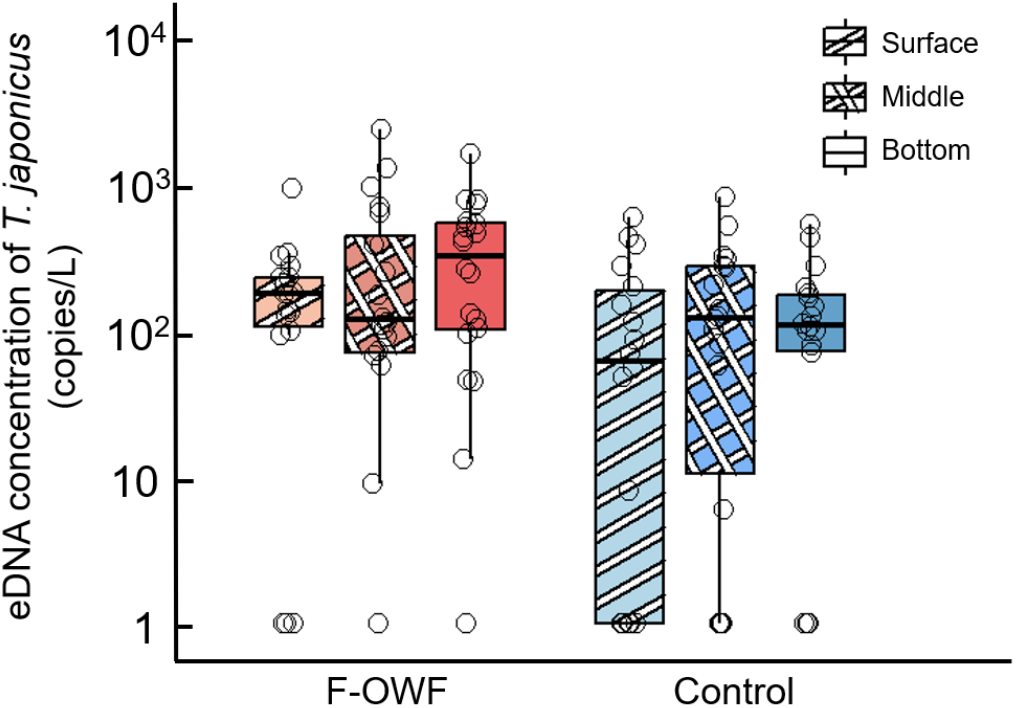
Comparison of eDNA concentrations between F-OWF and control stations across water layers (surface, middle, and bottom). The central line in whisker plots represents the median, whereas the lower and upper edges of the box indicate the first (25%) and third quartiles (75%), respectively. Whiskers represent minimum and maximum values, and individual data points are displayed as a scatter plot.

Since no significant differences were observed among water layers, Fig. 5 presents monthly mean differences in eDNA concentration using pooled data across water layers. In addition to sampling stations, sampling month also had a significant effect on eDNA concentration (*p* = 0.01, *p* = 5.33 × 10^−9^), while their interaction was not significant *(p* = 0.46) (Table 2), indicating that the relationship between F-OWF and control stations did not change with the season. The effect of sampling stations on eDNA concentration remained consistently positive throughout the year. Multiple comparisons among groups showed that eDNA concentration at F-OWF stations was highest in October, with significant differences compared to F-OWF and control stations in June and control stations in August (Fig. 5). In contrast, eDNA concentration at control stations was lowest in June, with significant differences compared to control stations in April, F-OWF and control stations in October, and F-OWF and control stations in December (Fig. 5).

**Table 2.** Summary of the two-way nonparametric ANOVA assessing effects of sampling stations (F-OWF and control) and month (Apr ~ Dec) on eDNA concentration. Sampling stations and months have a significant impact on eDNA concentration. In contrast, their interaction did not significantly affect eDNA concentration. Statistical significance is indicated as ** for *p* < 0.01 and * for *p* < 0.05.

| Factor | d.f | H statistic | p-value |
| --- | --- | --- | --- |
| Station<br>(Treatment/Control) | 1 | 5.92 | 0.01* |
| Month<br>(Apr/Jun/Aug/Oct/Dec) | 4 | 44.4 | $5.33 \times 10^{-9**}$ |
| Interaction<br>(Station × Month) | 4 | 3.60 | 0.46 |

**Fig. 5.**
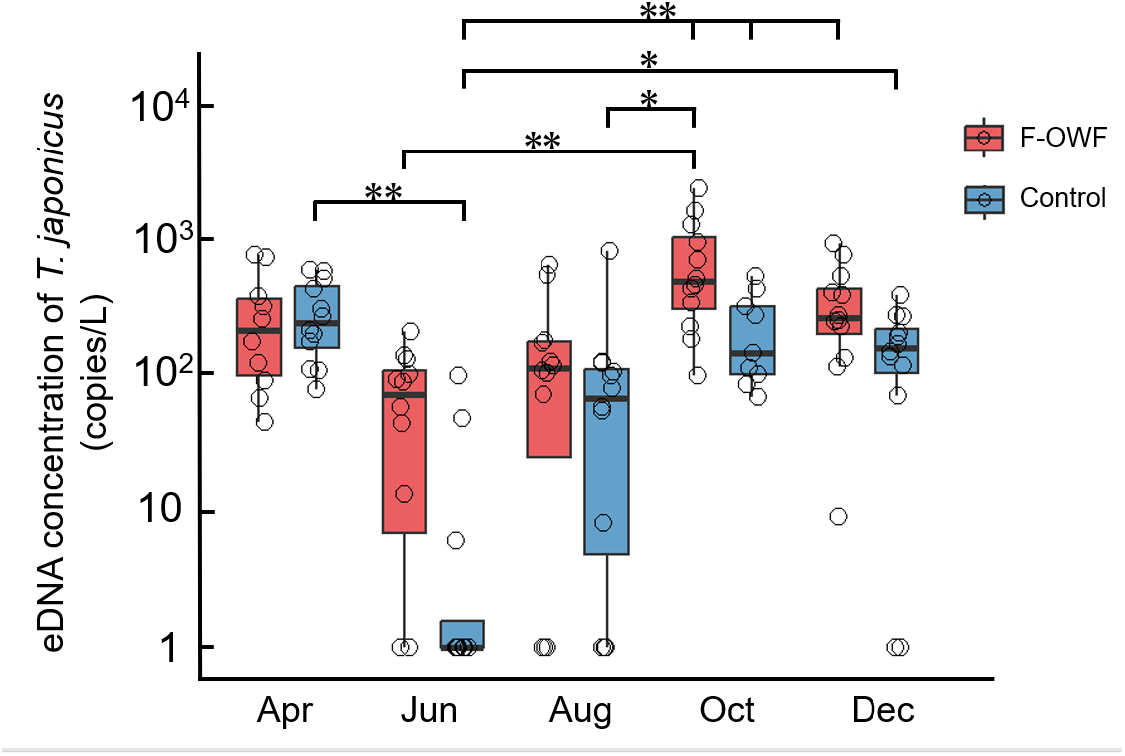
Comparison of eDNA concentrations between F-OWF and control stations and across months (April ~ December). The central line in whisker plots represents the median, whereas the lower and upper edges of the box indicate the first (25%) and third quartiles (75%), respectively. Whiskers represent minimum and maximum values, and individual data points are displayed as a scatter plot. Statistical significance is indicated as ** for *p* < 0.01 and * for *p* < 0.05.

Results of the GLM analysis showed that among environmental factors (presence of wind turbines, water layer (m), and water temperature (°C)), only wind turbines had a significant effect on eDNA concentration (*p* = 6.24 × 10^−3^) (Table 3). This indicates that eDNA concentrations derived from *T. japonicus* are influenced by the presence of wind turbines, regardless of other environmental factors.

**Table 3.** Summary of environmental factors affecting eDNA concentration based on the generalized linear model (GLM). The response variable was the log-transformed eDNA concentration, while explanatory variables included the presence of wind turbines and water layers. Statistical significance is indicated as ** for *p* < 0.01 and * for *p* < 0.05.

| Variable | Estimate | SE | t-value | p-value |
| --- | --- | --- | --- | --- |
| Intercept | 3.24 | 0.38 | 8.53 | $9.3 \times 10^{-14**}$ |
| F-OWFs<br>(Presence × Absence) | 1.12 | 0.40 | 2.79 | 0.01* |
| Depth (m) | 0.01 | 0.04 | 1.72 | 0.08 |

## DISCUSSION

Using eDNA technology, this study is the first to demonstrate the attraction of OWFs for fish, particularly *T. japonicus*. eDNA analysis is a powerful tool for collecting genetic information from organisms in aquatic or marine habitats, allowing researchers to study species distributions and biodiversity (Thomsen et al., 2015). Additionally, eDNA is released from biological sources such as excretions and tissues, and its concentration correlates with biomass (Doi et al., 2017). This suggests that eDNA concentrations are higher in areas where the target species is more abundant. However, since this study was conducted in an offshore environment rather than a closed system, careful consideration is required regarding how accurately eDNA reflects the actual biomass at sampling sites. One of the most important factors to consider is the potential persistence of eDNA. Fish-derived eDNA in marine environments decreases by 1.5–4.6% per hour, meaning that a large proportion of the detected eDNA is likely to have been released 1-2 days before sampling (Maruyama et al., 2014). However, an *in situ* eDNA experiment on *T. japonicus* conducted in Maizuru Bay, Japan, demonstrated that when the current velocity exceeds several tens of millimeters/s, eDNA disperses rapidly, further reducing the presence of older eDNA (Yamamoto et al., 2016). In this study, average current velocities at the sampling sites were 148 mm/s in April, 200 mm/s in June, 181 mm/s in August, 109 mm/s in October, and 141 mm/s in December. Therefore, the influence of eDNA persistence on biomass estimation in this study was likely minimal, and detected eDNA concentrations likely reflect the biomass of *T. japonicus* at the time of sampling.

In this study, eDNA concentrations of *T. japonicus* in the F-OWF stations near F-OWFs were, on average, significantly higher (105.13%) than at the control stations located farther from the F-OWFs (Figs. 2 and 3). Furthermore, GLM analysis of environmental factors affecting eDNA concentration indicated that the presence of F-OWFs was a significant explanatory variable, showing a positive effect on eDNA concentration (Table 3). Previous studies have reported that floating artificial structures can function as fish aggregation devices (FADs) (Castro et al., 2002; Inger et al., 2009). For example, Han et al. (2025) demonstrated that floating artificial reefs effectively attract epipelagic fish such as *Nibea albiflora*. Additionally, Okamoto (1992) reported that *T. japonicus* aggregates around floating reefs. The results of this study are consistent with these previous findings. Our findings suggest that F-OWFs off the western coast of Kyushu, Japan have a aggregating effect on *T. japonicus* and may function as FADs.

*Trachurus japonicus* may utilize F-OWFs throughout its life history. In this study, eDNA concentrations were consistently higher at stations near F-OWFs, regardless of the water layer (Table 1 and Fig. 4). The F-OWFs in the study area are installed at depths ranging from 100 to 135 m (Table S1) and consist of a floating section (submerged depth: 76 m) and chains extending to the seafloor, forming a vertically structured habitat. This suggests that *T. japonicus* is not aggregating in a specific layer, but rather in all layers around F-OWFs. One possible reason for this pattern is the influence of *T. japonicus* behavioral ecology at different growth stages. Juvenile *T. japonicus* form schools near the surface and gradually move to deeper waters as they grow (Enomoto et al., 2022; Nakamura and Hamano, 2009; Shida, 2002). In fact, adult fish have been collected using bottom trawl nets at depths of 100–200 m (Takahashi et al., 2012a). The floating structure of F-OWFs may provide shelter that mitigates wave and current impacts, attracting juvenile fish with limited swimming ability. Additionally, these structures may serve as refuges from predators. Indeed, studies have reported that species such as *Caranx crysos* aggregate around floating FADs to escape predation (Sinopoli et al., 2015). On the other hand, larger adults, which prefer rocky seabed habitats for shelter (Takahashi et al., 2012b), are likely to aggregate around the chain section of F-OWFs. For fish like *T. japonicus*, which transition between pelagic and demersal life stages, F-OWFs may provide an attractive habitat for *T. japonicus* throughout larval and adult stages. However, as this study did not involve direct capture or acoustic surveys, further investigations using acoustic depth-sounding technology and remotely operated vehicles (ROVs) are necessary to determine the actual residency patterns of *T. japonicus* around F-OWFs.

F-OWFs are likely to attract *T. japonicus* throughout the year. Although seasonal variations in eDNA concentration were observed, the influence of station (F-OWF and control) on eDNA concentration remained consistent throughout the year (Table 2 and Fig. 5). In waters off the western coast of Kyushu, Japan, *T. japonicus* reaches its peak spawning season in April, after which it is transported to the Sea of Japan by the Tsushima Warm Current over approximately 40 days (Kasai et al 2008; Sassa et al., 2009). In the southwestern Sea of Japan, juvenile *T. japonicus* abundance peaks along with the seasonal peak in prey availability (Fukataki, 1960). In autumn, as water temperatures decline, fish migrate southward, returning to waters off the western coast of Kyushu, Japan. This migration pattern suggests that *T. japonicus* that spawned in the western Kyushu move north in the spring, spend the summer in the Sea of Japan, and return south in autumn. Results of this study reflect this migratory pattern, as eDNA concentrations of *T. japonicus* decreased in summer and increased from autumn to spring (Fig. 5). This finding suggests not only an attraction of F-OWFs for fish, but also that the overall eDNA concentration may reflect the abundance of this species in this area. Notably, the effect of station type remained consistent throughout the year, indicating that eDNA concentrations appropriately reflect seasonal fluctuations in actual biomass. However, long-term effects of F-OWFs remain unclear. Long-term monitoring is necessary to evaluate whether F-OWFs contribute to establishment of fish schools and increases of local populations.

eDNA analysis alone cannot determine the size of individual *T. japonicus* congregating around F-OWFs. Although this study focused on a single species, a broader range of species must be examined to fully understand the impact of F-OWFs on fish communities. There are extremely few reports on the impact of F-OWFs on fish communities compared to those of B-OWFs, highlighting the need to expand the scope of species under investigation. Changes in dominant pelagic fish species from herring (*Clupea harengus*) to sand eels (*Ammodytes spp*.) have been reported around bottom-fixed offshore wind farms (B-OWFs) (Ybema et al., 2009). Other studies have documented significant increases in species such as common sole (*Solea solea*) and Whiting (*Merlangius merlangus*) near B-OWFs (Lindeboom et al., 2011). Furthermore, it has been estimated that fish availability around wind turbine foundations may be up to 50 times higher than in the surrounding sandy seabed (Leonhard and Pedersen, 2006). B-OWFs also provide refuge for cod (*Gadus morhua*) and pouting (*Trisopterus luscus*), affecting multiple reef-associated and demersal fish species (Stenberg et al., 2015). In addition, transmission cables used to deliver generated electricity emit electromagnetic fields, which may influence movements and navigation of species sensitive to electromagnetic and magnetic fields, particularly elasmobranchs, certain bony fish, decapod crustaceans, and sea turtles (Tricas et al., 2011; Gill et al., 2009; Westerberg et al., 2008). These findings indicate that OWFs likely impact a wide range of taxa. Migratory fish, in particular, tend to congregate around floating structures such as artificial reefs (Fréon & Dagorn, 2000). Studies have also suggested that combinations of bottom-fixed and floating artificial reefs can provide diverse habitats, attracting a wider variety of fish species (Han et al., 2025; Kellison et al., 1998; Wilhelmsson et al., 2006). The most significant difference between F-OWFs and B-OWFs is their deployment depth. While B-OWFs are restricted to shallower waters, F-OWFs can be installed even in deep-sea environments exceeding 200 m. If F-OWFs exhibit such reef effects, biomass changes distinct from those observed in B-OWFs may occur. Therefore, to effectively evaluate fish community composition and biomass changes, an integrated biological monitoring approach that combines traditional survey methods with metabarcoding eDNA technology is essential (Hinlo et al., 2017).

The study demonstrated that F-OWFs influence *T. japonicus* populations both spatially and temporally, but is this necessarily beneficial for fishermen and fish predators? The answer is not simple. Construction of F-OWFs leads to establishment of no-fishing zones or refugia (Reckhaus, 2022). If fish congregate in these restricted areas, fishermen may experience negative impacts on their catches. Additionally, since fishermen must navigate around F-OWFs, they must use extra fuel to avoid them (Nakao, 2022). There is also the possibility that fish that would have otherwise entered set nets may instead aggregate around F-OWFs, reducing catch efficiency. The era has arrived in which fish must coexist with offshore renewable energy generation. Moreover, F-OWFs not only impact fish populations; they also pose potential problems for humans, such as noise pollution, and for birds, in the form of bird strikes (Furness, 2013). A large-scale monitoring program should be implemented and integrated with adaptive development of future wind farms. This monitoring should be conducted continuously across all phases—before, during, and after construction. Furthermore, it is essential to investigate not only operational impacts of wind farms, but also the intensity and effects of other human activities in marine environments (Lindeboom et al., 2011).

## Acknowledgments

We sincerely thank the crew of the training vessel (*T/S*) Kakuyo-Maru for their great support in this research. We also appreciate the staff of the Fish and Ships Laboratory, Faculty of Fisheries, Nagasaki University. Finally, we gratefully acknowledge the financial support from the Nissui Life Foundation for this research.

## Author Contributions

ST Investigation, Writing – original draft, Visualization. RK Investigation, Writing – review & editing. SN Investigation, Writing – review & editing. SA Investigation, Writing – review & editing. SSB A Investigation, Writing – review & editing. ST Investigation, Writing – review & editing. SM Investigation, Writing – review & editing. JU Investigation, Writing – review & editing. TA Investigation, Writing – review & editing. KH Investigation, Writing – review & editing. SF Investigation, Writing – review & editing. KS Investigation, Writing – review & editing. MH Investigation, Writing – review & editing. MY Conceptualization, Methodology, Investigation, Writing – original draft, Supervision, Funding acquisition, Visualization.

## Supplementary Materials

**Table S1.**
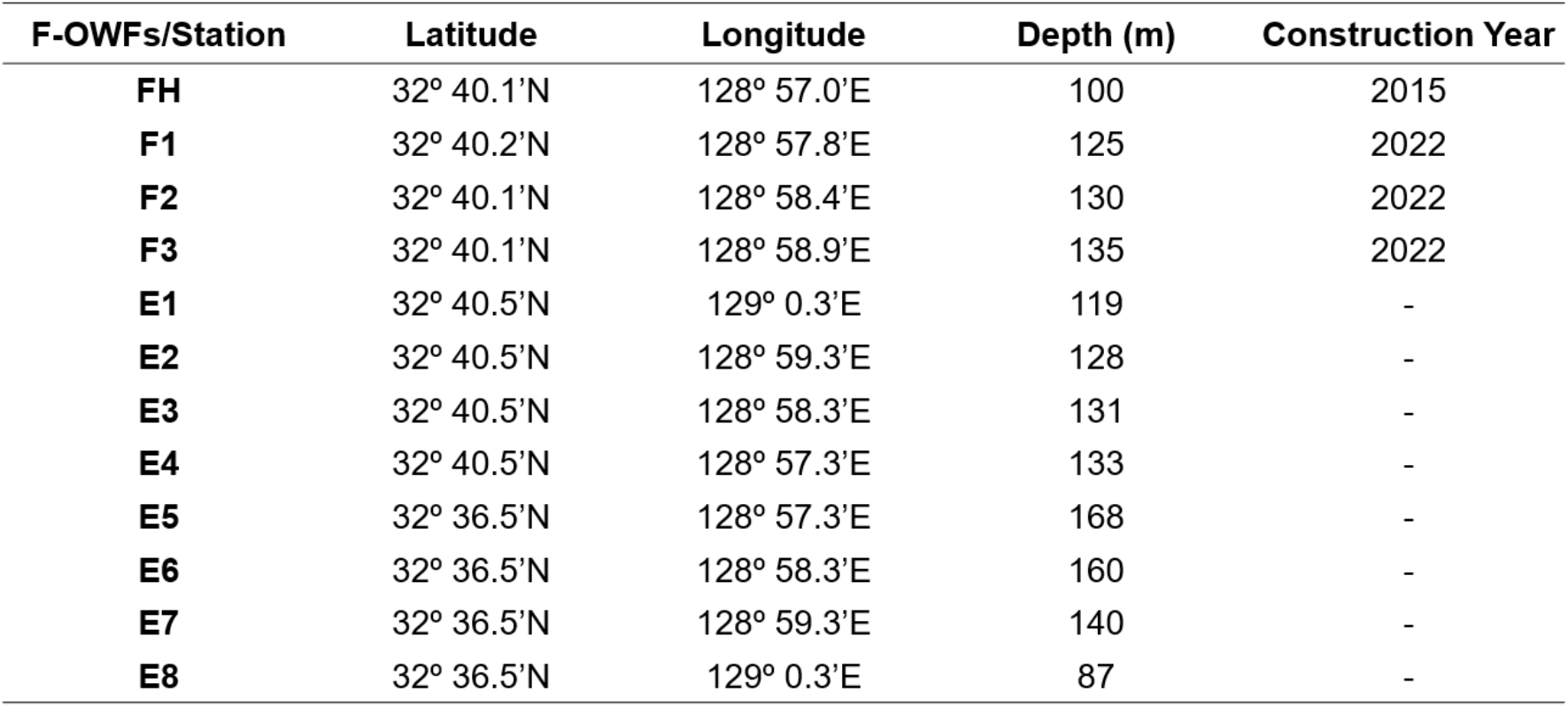
Latitude, longitude, and depth of each station and F-OWF locations (FH, F1, F2, F3). Construction dates of F-OWFs are also documented, and locations of each site are shown in Fig. 2.

| F-OWFs/Station | Latitude | Longitude | Depth (m) | Construction Year |
| --- | --- | --- | --- | --- |
| **FH** | 32° 40.1'N | 128° 57.0'E | 100 | 2015 |
| **F1** | 32° 40.2'N | 128° 57.8'E | 125 | 2022 |
| **F2** | 32° 40.1'N | 128° 58.4'E | 130 | 2022 |
| **F3** | 32° 40.1'N | 128° 58.9'E | 135 | 2022 |
| **E1** | 32° 40.5'N | 129° 0.3'E | 119 | - |
| **E2** | 32° 40.5'N | 128° 59.3'E | 128 | - |
| **E3** | 32° 40.5'N | 128° 58.3'E | 131 | - |
| **E4** | 32° 40.5'N | 128° 57.3'E | 133 | - |
| **E5** | 32° 36.5'N | 128° 57.3'E | 168 | - |
| **E6** | 32° 36.5'N | 128° 58.3'E | 160 | - |
| **E7** | 32° 36.5'N | 128° 59.3'E | 140 | - |
| **E8** | 32° 36.5'N | 129° 0.3'E | 87 | - |

